# SVlog: a logic programming framework for understanding structural variation in genomic disease

**DOI:** 10.64898/2026.08.11.744322

**Authors:** Mikhail Gudkov, Andre L. M. Reis, Meutia Kumaheri, Ira W. Deveson

## Abstract

Structural variants (SVs) are a diverse group of genetic variants defined by a minimum size of 50 base pairs. SVs account for the majority of all variant bases in a person’s genome and are commonly implicated in inherited disease and cancer. However, SV analysis is complex due to their wide variation in type and size, degree of polymorphism, involvement of repetitive sequences, and the myriad ways they may elicit a functional impact, as well as technical factors like imprecise breakpoint detection, and alternative representations of the same event. Despite recent advances in the detection and characterisation of SVs, it remains difficult to assess them beyond basic annotations and comparisons.

Here we introduce SVlog, a transparent and extensible meta-programming framework for SV analysis. With the logic programming language Soufflé as its engine, SVlog provides a declarative ontology describing relationships among SVs, genes and other genomic elements. Genome annotations and SV datasets – both user-provided and public reference data – are converted into relational ‘facts’, to which SVlog applies logical rules that define ‘predicates’. Predicates are specific, transparent and deterministic, yet fully flexible and composable, enabling detailed evaluation of SVs without relying on stochastic “black box” approaches.

To showcase SVlog, we have developed a ready-made predicate library for SV annotation, comparison and prioritisation in the context of rare inherited disease. Despite its compact codebase, SVlog evaluates more than 50 input predicates to generate over 70 informative output predicates. It synthesises evidence from population and clinical genomic databases, and applies a tiered filtering strategy to identify candidate pathogenic SVs in patients with inherited disease.

By focusing on explainability and modularity, SVlog offers a fast, reliable library for SV analysis and is a powerful deterministic alternative to traditional bioinformatics pipelines for clinical variant curation.

## Introduction

Rare inherited disease studies are often hindered by the amount of data associated with each genetic variant and difficulties in using that data in a systematic way to find true causal variants. This is particularly the case when using long-read sequencing technologies, which, despite increasing the diagnostic yield overall, tend to generate larger amounts of data by resolving variants that cannot be detected with short reads (Del Gobbo and Boycott, 2025; Steyaert *et al*., 2025).

One class of genetic variation that benefits from the advances in genetic sequencing is structural variants (SVs) (Collins *et al*., 2020). Accounting for the vast majority of varying base pairs between any two human genomes, SVs play an important role in human genomics. However, even though the detection of structural variants has improved substantially over the last decade, the analyses associated with this type of variation — such as clinical variant prioritisation — lag far behind.

Unlike single nucleotide variants (SNVs), SV interpretation presents unique computational and biological challenges: they encompass a vast diversity of event types that vary in size, disproportionately occur in repetitive sequence contexts, and are highly polymorphic (more so than SNVs). In addition, the identification of SV breakpoints is often imprecise due to technical and analytical limitations, and it may be valid to represent a given SV in multiple different configurations. These factors confound the analysis of SVs, whether tracing pathogenic drivers in rare inherited diseases, disentangling structural rearrangements in cancer, or cataloguing allele frequencies in population genetics. Long-read sequencing technologies, alongside better calling algorithms, tend to resolve SVs with greater sensitivity and precision and have, therefore, become crucial in genomic disease studies (Jensen *et al*., 2025). To our knowledge there is no widely adopted framework for representing arbitrary relations between SVs (Kirsche *et al*., 2023). Hence, the problem of comparing SVs still relies on *ad hoc* solutions.

The interpretation of genetic variants in rare inherited disease follows a triage workflow that narrows the tens of thousands of variants typically called per genome down to a handful of candidate drivers. The lines of evidence used in this triage include allele frequency in reference populations (common variants may be assumed to be benign), predicted impact on coding and regulatory sequence elements, gene-level metrics of sequence constraint (e.g. LOEUF (Karczewski *et al*., 2020)), disease relevance of impacted genes or elements (e.g. based on OMIM and curated panels such as the Mendeliome), clinical assertions in databases such as ClinVar (Landrum *et al*., 2020), and inheritance patterns from family genotypes. These criteria are codified in standards such as the ACMG/AMP guidelines, and clinical interpretation tools combine them through composite scores or rule-based filtering.

This workflow has yet to be effectively adapted for SVs. Population reference data for SVs has only recently approached the depth available for SNVs, and the largest datasets (e.g. gnomAD-SV v4.1) are inherently limited by their use of short-read sequencing data. SVs are also sparsely annotated in ClinVar and other clinical databases. Meanwhile, several lines of evidence used for SNVs have no direct SV counterpart — for example, there is no standard equivalent of a CADD or AlphaMissense score for an arbitrary SV. Existing tools such as AnnotSV (Geoffroy *et al*., 2018) and SVAnnotate (Collins *et al*., 2020) provide annotation and prioritisation pipelines, but they encode fixed rule sets and scoring schemes that are difficult for the user to inspect, modify or extend. There remains a gap for systems in which the prioritisation logic is itself the object of work – readable, editable, and adaptable to the cohort, gene panel or disease context at hand.

In the last few years there has been a proliferation of machine learning (ML) methods aimed at converting genomic data into models that can predict which variants are likely to be deleterious and therefore would be good candidates in disease studies (Jensen *et al*., 2025; Kleinert and Kircher, 2022; Cheng *et al*., 2023; Parry *et al*., 2024). However, most advanced ML methods can be seen as ‘black boxes’ that make predictions in ways that are not directly obvious to the user (Quinn *et al*., 2022; Van Smeden *et al*., 2022). Despite explainable Artificial Intelligence (XAI) methods becoming more popular for those same reasons (Liu and Xu, 2023; Farahani *et al*., 2022; Vassiliades *et al*., 2021), we believe that many applications that they are used for do not require the level of complexity that comes with those methods. With the increasing profile of complex ML/AI strategies like neural networks and large language models, there is a risk of overlooking simpler, deterministic and transparent solutions.

One such method is *logic programming*, a declarative programming paradigm where the user defines facts and rules rather than step-by-step instructions. So far, applications of logic programming in the life sciences have been limited to the modelling of biological networks (such as those describing cell signalling and metabolic processes) (Hall and Niarakis, 2021; Wynn *et al*., 2012; Hemedan *et al*., 2022) and integration of biological databases (Gupta *et al*., 2010; Angelopoulos and Giamas, 2015). A particularly interesting example of how logic programming helps to address challenges in understanding biological mechanisms of diseases is cancer pathways (Palma *et al*., 2021; Béal *et al*., 2021, 2019) and disease maps in general (Ostaszewski *et al*., 2019). Despite the fact that there is relatively little interest in logic programming from the medical community, these methods continue to evolve and become more capable. For example, an extension of general logic programming methods called Answer Set Programming (ASP) has also seen some applications in biology (Frioux *et al*., 2020).

We, therefore, hypothesised that logic programming would provide a perfect trade-off between minimalism and expressivity in addressing the challenges associated with genomic SV analysis. In our efforts to systematise all relevant pieces of information relating to SVs, we created SVlog, a logic programming library containing a set of ready-made predicates for SV analysis in the context of rare inherited disease. The SVlog library is designed to establish relationships between SVs detected in a patient/family and SVs detected in other members of a research cohort, population reference databases, and genomic elements including genes, repeats and regulatory elements. This addresses an immediate capability gap for the field, and establishes SVlog as an efficient, deterministic and extensible ontology for SV analysis.

## Methods

The methodology developed here is based on the principles of logic programming (Figure 1). Specifically, we chose Soufflé, a dialect of Datalog, as our modelling language (Jordan *et al*., 2016). Datalog can be seen as a minimal, restricted subset of Prolog: even though Datalog is not Turing-complete like Prolog, it is a fully declarative language, unlike Prolog, which is not strictly declarative. Over the last decade, Soufflé has established itself as a widely used, high-performance Datalog engine. In this work, we used Soufflé version 2.5.

**Figure 1.**
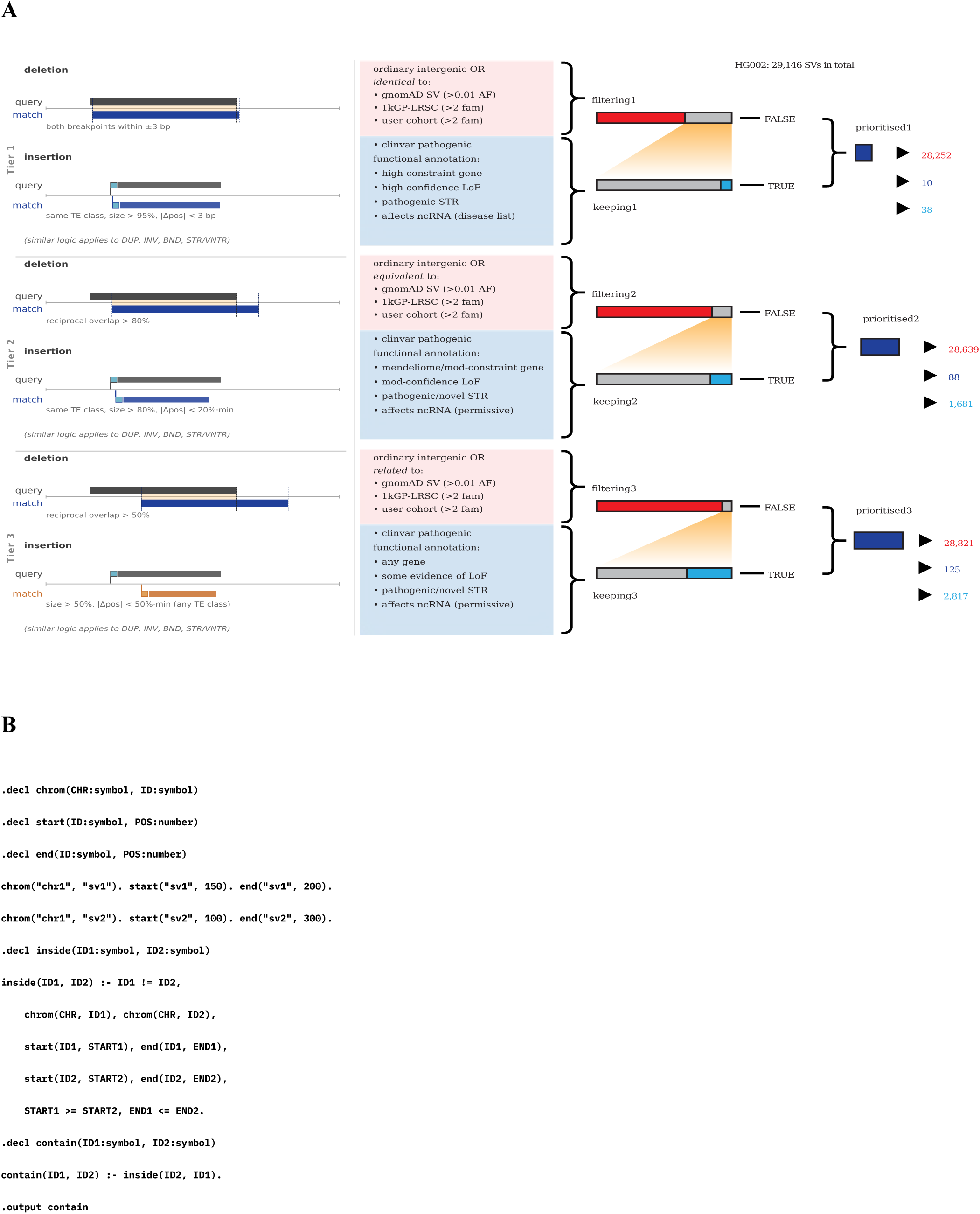
Implementation of the variation prioritisation strategy in SVlog. (A) The tiered prioritisation logic. (B) Minimal working example of a predicate implemented in SVlog.

In our framework, each genomic element (variants, exons, TADs, etc.) comes from a dataset and has a unique ID. Each dataset, which is usually a VCF file, is processed in such a way that annotations for each element are converted into so-called “facts”. For example, variants are defined using such predicates as `chrom()`, `start()`, `len()` and `type()`. Facts look like this: ‘`chrom(“chr1”, “var1”).`’. In Soufflé, facts are usually a collection of TSV files, where each column is a positional argument in the corresponding predicate. For example, a TSV file ‘chrom.facts’ containing ‘chr1 var1’ can be used to indicate that a variant with ID ‘var1’ comes from chromosome 1 (‘chr1’), in the same way as above.

Facts are the input data (Extensional Database, EDB) for rules, which, in turn, define possible relations between genomic elements based on some criteria. For example, a rule determining whether two variants overlap would need to use the information from each variant’s `chrom()`, `start()` and `len()` facts.

Rules can be nested: complex rules can be defined using several simpler ones. Composability is one of the main features of Datalog. In a way, individual predicates in Datalog can be seen as LEGO pieces forming blocks. Datalog is a declarative language, which means every individual block is independent of the other blocks and addresses a specific part of the problem that is defined in its natural context. Soufflé uses a semi-naive bottom-up evaluation strategy, which means that to find all sets of values for which a predicate evaluates to “true”, Soufflé will evaluate all intermediate predicates that the target predicate depends on. For many applications, this is not practical, as intermediate predicates often define general cases, which, when evaluated in full on large amounts of data, require substantial memory allocations. An improvement over the semi-naive bottom-up strategy is the Magic Set Transformation. This transformation re-writes the original Datalog program in a way that emulates top-down evaluation, which is similar to how Prolog generally operates. Magic Sets, therefore, can direct the Datalog engine to compute only those facts that are necessary to generate a complete set of output facts (Intensional Database, IDB).

SVlog analyses all input SVs and does not require FILTER=PASS, because its annotation is orthogonal to caller QC and we wanted to test it across the widest set of SVs (including low-confidence ones), leaving de-prioritisation to the user. Where possible, SVs were annotated for repeat and mobile-element context with our companion tool SVscanner (https://github.com/GenTechGp/SVscanner).

### Data

Several datasets were converted into fact tables for our framework, namely gnomAD (version 4.1), ClinVar (version “2024-11-11”), an in-house cohort of 485 individuals from 184 families (199 probands), as well as 264 samples from Oxford Nanopore long-read resequencing of 1000 Genomes Project samples, processed using the same pipeline as the in-house cohort (Figure 2). We also used GENCODE version 46 (Frankish *et al*., 2023) for reference gene elements (canonical transcripts from protein-coding genes only). Additionally, we used separate data for regulatory elements (GeneHancer, RefSeq), repetitive elements (STRchive, STRipy, Tandem Repeat Finder), gene constraint levels (LOEUF), gene panels (OMIM, Mendeliome) and noncoding RNA lists (seqr).

**Figure 2.**
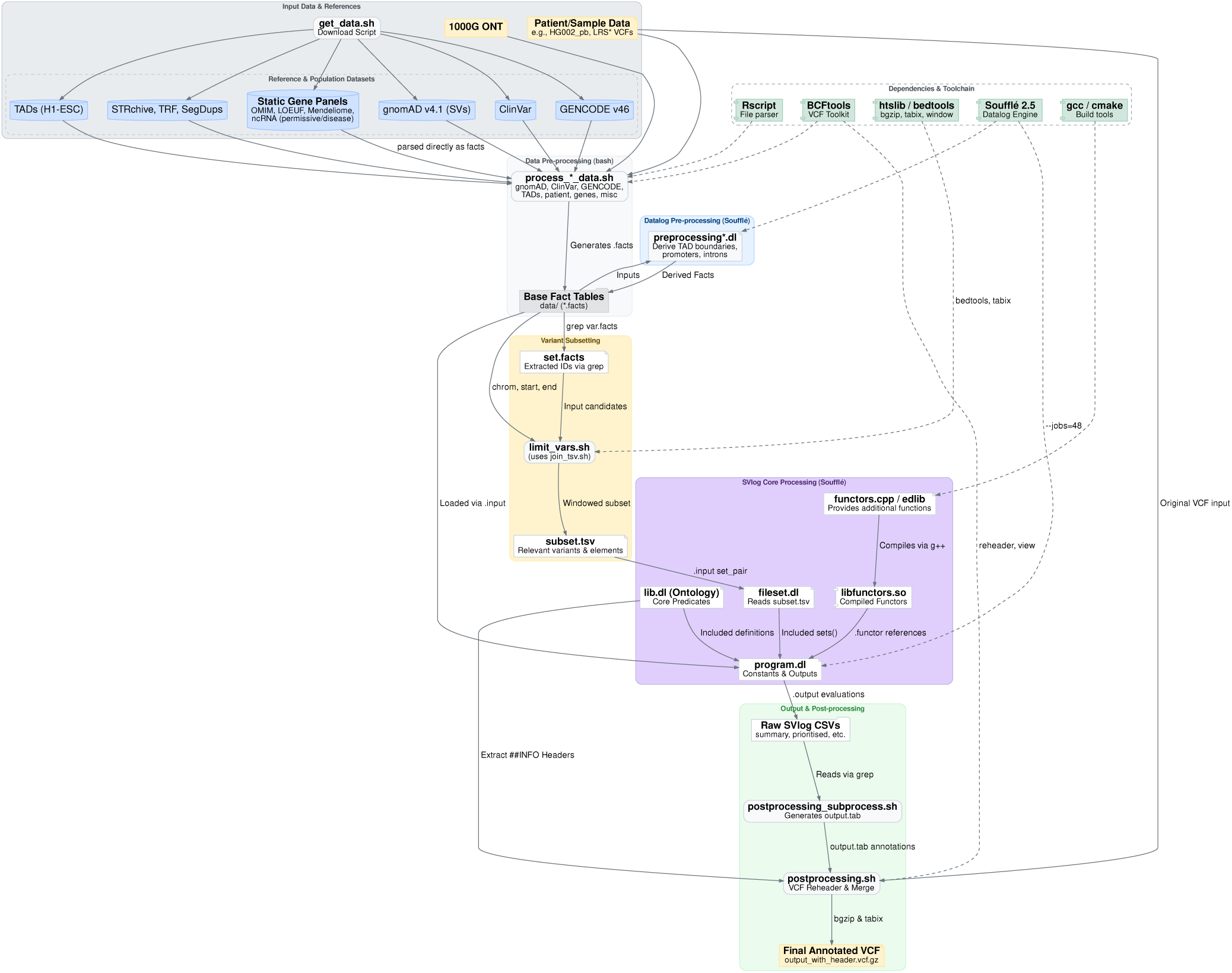
Data workflow in SVlog. Reference data is first downloaded and converted into “facts”, which are then augmented with additional information derived with Soufflé; target variants are extracted and pre-processed with BEDtools to generate a list of matches within each variant’s window, to limit the computational burden; the generated pairs are then processed through SVlog and the original VCF is annotated with the derived predicates.

## Results

### SVlog

SVlog is an ontology of events in the domain of genomic structural variation, built on a logic programming framework. A structural variant (SV) or other genomic element (e.g. genes, enhancers, etc) is represented as a set of “facts”, which include its genomic coordinates, element type, and other descriptive information.

Facts are stated relative to simple predicates, such as `chrom()`, `start()`, `len()`, `type()`. For example, predicates and facts pertaining to an SV (`“var1”`) found in the ClinVar database might include `‘chrom(“chr1”, “var1”)’`, `‘type(“deletion”, “var1”)’`, `‘from(“clinvar”, “var1”)’`, `‘clnsig(“Pathogenic”, “var1”)’`. Meanwhile, predicates and facts pertaining to an evolutionarily constrained gene (’gene1’) might include `‘omim(“gene1”)’` and `‘constrained_gene(”gene1”)’`. Predicates may be nested and connected by logical operators (’AND’, ‘OR’, ‘EQUAL TO’, etc), allowing complex rules to be constructed using multiple simpler ones. Within the SVlog ontology, every genomic element is thus described by a flexible and open-ended set of predicates and facts.

Predicates may be written to describe relationships between two elements, such as the relationship of an SV to a gene or to another SV. Whereas two single nucleotide variants (SNVs) from different sources can be unambiguously matched using their genomic coordinates and base substitutions, the comparison of two SVs requires more nuance. Two SVs do not need to be *identical* to have the same functional impact. Depending on the context, it may be more useful to understand whether they are *equivalent* or even *related*, where these comparative terms consider the two SVs’ types, sizes, degree of overlap, repeat context (e.g. repeat expansion, mobile element insertion, etc) and their relationships to other genomic elements (e.g. impact on a gene or enhancers). The SVlog core predicate library therefore includes the hierarchical predicates `identical_variants()`, `equivalent_variants()` and `related_variants()` to describe the similarity of any two SVs (Figure S2). The relationship of an SV to a gene or other genomic element is similarly contextual. The potential for an SV to impact the function of a protein-coding gene depends on its type, size, orientation, sequence context and position within the gene’s architecture. The SVlog library contains simple predicates like `affects_cds()` or `affects_only_promoter()` to encode these parameters, as well as more complex predicates predicting loss-of-function (LoF) gene effects of various types (Table S1). The efficient evaluation of relationships among many SVs and genomic elements from different sources, according to nuanced yet flexible comparison criteria, is one of SVlog’s key capabilities.

SVlog may also be understood as a computer library containing definitions of many logical predicates relating to SVs. In conjunction with the Soufflé engine (Jordan *et al*., 2016) (see **Methods**), this library may be used to annotate SVs or answer specific questions about their biology. SVlog is a flexible system for exploring SVs at the level of the individual, family, cohort or population, where predicates can be written for any research question. One immediate use-case is for the interpretation of potential pathogenicity of SVs in the context of inherited disease. We have, therefore, developed a basic, ready-to-use predicate library for this task, explored in detail in the following sections. Despite the density of inter-predicate relations (Figure S1), this library is highly compact. With comments and empty lines removed, the library contains just under 1, 600 lines and 50, 000 characters, of which >20% are directives specific to Soufflé. Such brevity is enabled by Datalog’s inherent algorithm-free, structureless nature that does not require any programming *per se* (Jordan *et al*., 2016) (Figure 1B). The current latest version of SVlog (0.99.146) processes information from over 50 input predicates and outputs over 70 “conclusion” predicates useful for downstream SV analysis (see Table S1). In our view, it would be very difficult to implement code encompassing the full complexity of this predicate library using conventional programming languages. The evaluation of input data against this predicate library is computationally efficient, processing a typical human SV callset containing 26, 000 SVs against a large set of population reference data and genome annotation files (see below) in just under 14 hours, using 48 CPUs and at least 130 GB of RAM, with all conclusion predicates evaluated and their associated facts annotated on the original VCF file. Importantly, these compute requirements were only necessary because of the large volume of input data and reference data, and can be more sparing for smaller input datasets.

### Prioritising variants with SVlog

The SVlog library relates each proband SV to variants in a research cohort and in population reference databases, and to genomic elements including genes, repeats and regulatory elements, so as to inform the user of the potential pathogenicity of a given SV. The logic, structure, annotations and datasets used are outlined in Figures 1 and 2, and described below. Table S1 contains a list of major predicates in the current library.

Every human genome harbours tens of thousands of SVs, with whole-genome long-read sequencing (LRS) typically detecting >20, 000 SVs per individual (Del Gobbo and Boycott, 2025). When considering a patient with a rare inherited condition, the diagnostic challenge lies in filtering through this complex background to find one or more candidate pathogenic variants. The pool must be narrowed down to SVs that are rare in the general population, are predicted to disrupt genes relevant to the patient’s clinical phenotype, and appropriately segregate with the condition based on family inheritance patterns (Jensen *et al*., 2025).

SVlog allows us to automate and standardise this reasoning. To establish rarity, SVlog ingests population allele frequencies from gnomAD (v4.1) and the 1000 Genomes Project Long-read Sequencing Consortium (1KGP-LRSC), clinical assertions from ClinVar, and, optionally, variants detected in a user’s own research cohort. In our analysis below, we utilise an in-house cohort comprising 184 families sequenced with LRS to improve variant filtering. A user may substitute this for their own cohort, as relevant to their research context, or simply proceed using only publicly available reference data. To evaluate genomic context, SVlog integrates reference annotations from GENCODE (transcripts, exons, UTRs), regulatory topologies (enhancers, TAD boundaries), and repetitive element databases (STRchive, STRipy, and SegDups). Finally, clinical relevance is weighted using established gene constraint metrics (LOEUF) and clinical gene panels, including OMIM, the Mendeliome, and curated lists of noncoding RNAs.

SVlog allows researchers to interact with this volume of data in two primary ways. Given a single candidate SV of interest, SVlog can rapidly retrieve all its associated information: identifying equivalent or related variants, evaluating their population frequencies, predicted impact on genomic elements, and relevant clinical annotations. Alternatively, SVlog can agnostically evaluate the entire set of SVs found in a single proband, autonomously filtering and prioritising SVs according to deterministic rules in the predicate library. All SVs are labelled with descriptive output predicates and their associated facts; however, the framework is designed to quickly identify the most relevant candidates, which meet the criteria for one or more of the apex predicates `prioritised1()`, `prioritised2()` or `prioritised3()`. These are underpinned by hierarchical rules that serve to prioritise (“keeping” predicates) or de-prioritise (“filtering out” predicates) a given SV, themselves drawing on lower-level facts about its rarity, gene impact, etc.

The ‘keeping’ predicates assimilate evidence *in favour* of the potential pathogenicity of a proband SV, including predicted LoF gene effects and relatedness to previously known pathogenic variants. LoF predicates are granular and tailored to specific variant types, such as `lof_translocation()` (BND records with REF position in or close to a gene) and `lof_strong()` (SV affects CDS, is splice-altering or is a within-gene translocation). These feed into higher-level prioritisation rules that weigh up both the strength of the LoF prediction and the functional relevance of a given gene, drawing on three complementary sources of gene-level evidence. LOEUF provides a continuous per-gene score of intolerance to loss-of-function variation (Karczewski *et al*., 2020). SVlog discretises it into two thresholds, flagging genes in the top 20% of constraint as “highly constrained” and the top 50% as “moderately constrained”. The Mendeliome is a broad, curated panel of ∼5, 000 genes with established or emerging Mendelian disease associations, maintained on PanelApp (Martin *et al*., 2019) and used here as a generic disease-relevance filter for probands without a specific phenotype hypothesis; only genes rated “green” (high-confidence, expert-reviewed) are treated as panel members. When a genotype-phenotype hypothesis exists, the user can supply any PanelApp panel via the `my_panel()` input, which SVlog treats identically to the Mendeliome but restricted to the relevant gene set. These three signals combine with the LoF categories to define graded prioritisation predicates: `prioritise_lof_high` (LoF-strong in a top-20% LOEUF gene), `prioritise_lof_mod` (LoF-strong or LoF-moderate in a top-50% LOEUF gene), `prioritise_lof_mendeliome` (LoF-strong or LoF-moderate in a green Mendeliome gene), and `prioritise_lof_my_panel` (LoF-strong or LoF-moderate in a user-panel gene), with `prioritise_lof_any` retained as a permissive fallback across any gene. For more complex variant types, SVlog combines multiple data streams into more specific queries. For example, to identify pathogenic short tandem repeats (STRs), the engine checks if an insertion variant is adjacent to a known pathogenic STR site and evaluates whether the insertion length represents a significant expansion. A proband SV may trigger several of the above predicates simultaneously, and they are together used to rank candidates into groups with decreasing evidence of pathogenicity: `keeping0()`, `keeping1()`, `keeping2()`, `keeping3()`.

The “filtering out” predicates assimilate evidence *against* the potential pathogenicity of an SV, based on comparison to the variation catalogs and annotations listed above. The variation catalogs in current use have complementary strengths and limitations. gnomAD-SV (v4.1) is the largest publicly available and openly downloadable SV reference, providing allele frequencies from short-read sequencing across diverse populations. Its scale makes it well suited for setting common-variant thresholds, but short-read calling misses a substantial fraction of the SVs recovered by LRS, particularly in repetitive and low-complexity regions. 1KGP-LRSC (264 samples here) is an LRS counterpart processed through the same Sniffles2-based pipeline (Smolka *et al*., 2024) as our in-house cohort. It captures SVs that gnomAD-SV does not, but encompasses many fewer individuals, and per-site frequency estimates are noisier as a result. ClinVar (release 2024-11-11) is curated rather than population-based, contributing positive and negative controls (pathogenic and benign assertions) used for both prioritisation and filtering, although SV coverage is sparser and less uniform than for SNVs. Our in-house cohort comprises whole-genome LRS data on 485 individuals from 184 families with 199 probands, processed through the same workflow and analysis pipeline as any proband sequenced in our laboratory. This should be understood as a user-provided cohort, generated from whatever suitable SV calls are available locally. While there is value in utilising local data, a user cohort is not required to run SVlog: filtering can operate on public datasets alone. A proband SV is de-prioritised if it is linked to a common SV in any of these catalogs via the `identical_variants()`, `equivalent_variants()` and `related_variants()` predicates described above. The `ordinary_intergenic()` predicate is also used to de-prioritise proband SVs with no proximity to genes or regulatory elements. These predicates feed into a hierarchy of predicates with increasing evidence against pathogenicity: `filtering_out1()`, `filtering_out2()`, `filtering_out3()`.

The evidence both for and against an SV’s pathogenicity is hence organised in tiers based on the degree of similarity to annotated variants and the strength of functional and clinical evidence. This lets the user relax the prioritisation criteria when the top tier yields no good candidates, while still ranking candidates: those surviving the strictest tier are the strongest, and looser tiers surface progressively weaker but still informative signals. Importantly, the tiers are not strictly nested: because a more permissive tier matches more variants and therefore also filters more out, a candidate prioritised under a stricter tier can be filtered out under a looser one (so the set of SVs prioritised across all tiers is larger than the set from any single tier). Ultimately, SVlog intersects these predicates and an SV is prioritised if it safely bypasses the “filtering out” exclusion rules while triggering one or more “keeping” conditions. Once again, these rules are tiered: `prioritised1()` applies the strictest thresholds to both filtering and keeping, `prioritised2()` relaxes both moderately, and `prioritised3()` is the most permissive on both axes. The tiers differ first in the comparison used to pair a proband SV with catalogue SVs — `identical_variants()` in Tier 1, `equivalent_variants()` in Tier 2 and `related_variants()` in Tier 3 — corresponding to progressively more permissive matching (Figure S2). Finally, when sequencing data from additional family members is available, SVlog may automatically establish inheritance patterns to further support variant interpretation. For example, the `denovo()` predicate tags any SV that is present in the proband but strictly absent in both sequenced parents. Overall, the predicate library is an efficient, nuanced and ready-made framework for SV prioritisation in the context of rare inherited disease.

### Analysis of HG002 reference sample

To evaluate this framework, we analysed LRS data from the popular human genome reference sample HG002. The sample was sequenced using both Oxford Nanopore Technologies (ONT) and PacBio whole-genome LRS and processed through an open source analysis pipeline developed by our team (https://github.com/leahkemp/pipeface), producing 29, 146 and 27, 557 SVs, respectively. These SV callsets were separately evaluated with SVlog, using the inherited disease predicate library described above, and taking the HG002 individual as a mock proband (noting the HG002 donor individual was not known to be affected by a rare inherited disease). The results for the ONT dataset are presented in Figures 3 and 4, while the PacBio data is presented in Figure S3.

**Figure 3.**
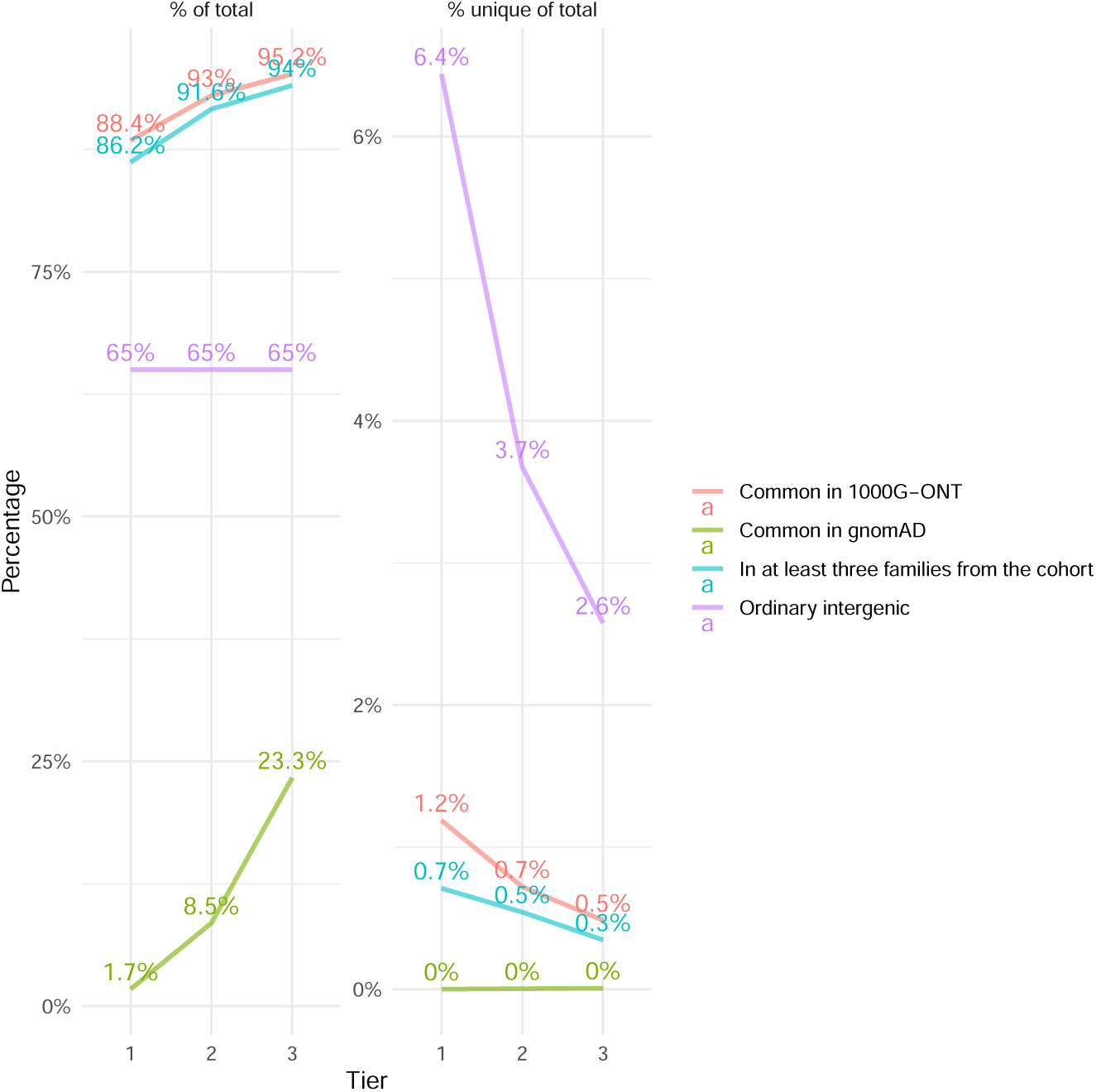
Performance of the filtering categories and tiers. HG002-ONT data (29, 146 variants). The left pane shows the percentage of variants matching each filtering set of conditions for filtering out; the right pane shows the percentage of variants that would be left unfiltered in the absence of the corresponding set of filtering conditions. As the stringency of comparisons decreases from Tier 1 to Tier 3, more variants start matching other variants in the database, which results in a higher percentage of filtered-out variants and a greater degree of sharedness between the filtering categories. The “Ordinary intergenic” category does not rely on other variants, hence there is no change in the rates when comparing the three tiers.

**Figure 4.**
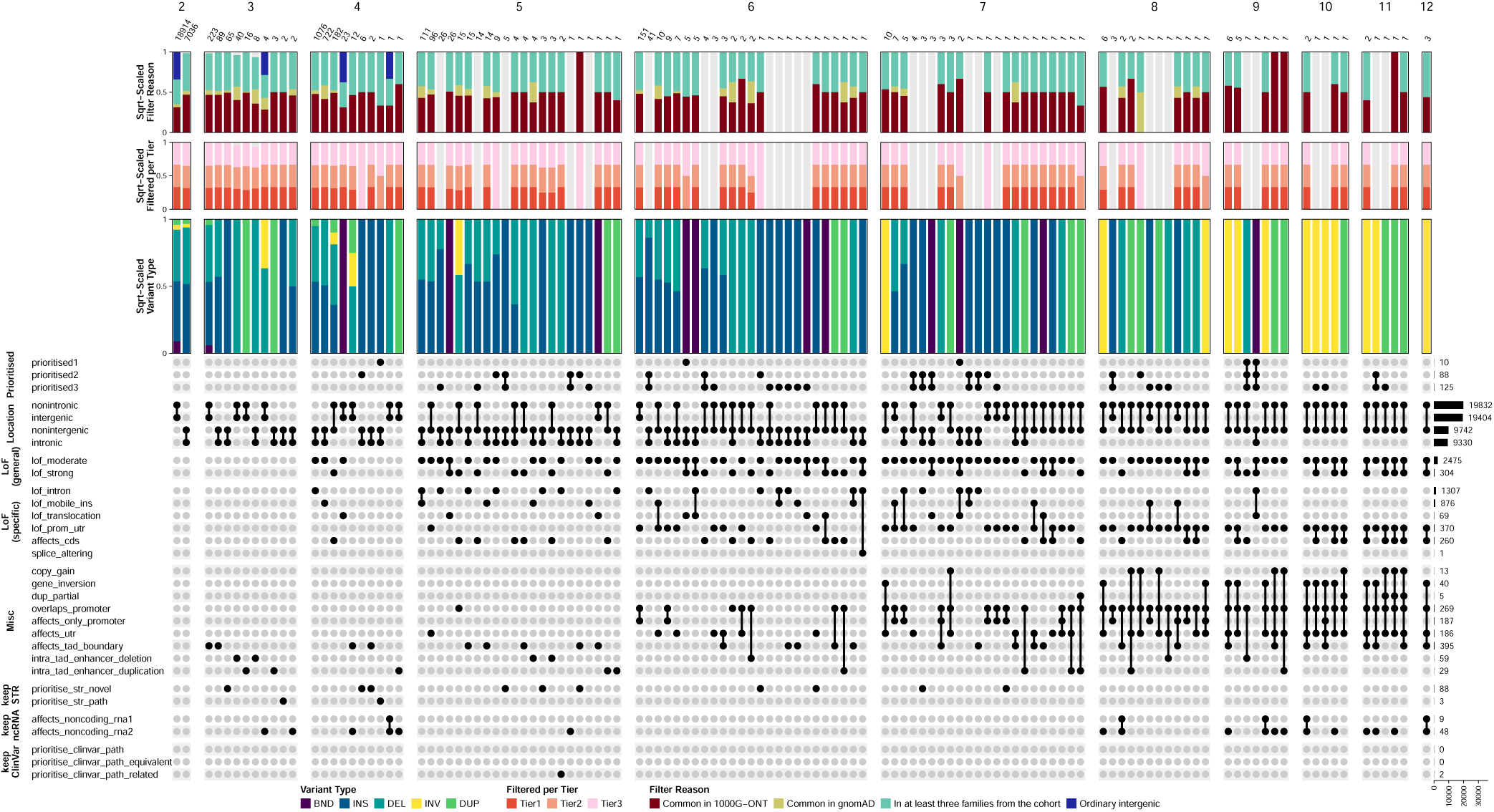
Prioritisation and variant effect predicates (rows) and predicate signatures (columns). HG002-ONT data (29, 146 variants). Only the most relevant predicates are included. Selectivity of each predicate is shown on the right. The panes are split based on the number of predicates in each signature.

First considering SV calls from ONT data, we assessed the performance of SVlog’s “filtering out” predicates relative to the different variation catalogs that inform filtering. Of the 29, 146 input SVs, the `filtering_out1()`, `filtering_out2()`, `filtering_out3()` predicates were triggered by 97% (28, 252), 98% (28, 639) and 99% (28, 821) of SVs, respectively. Parsing by reference data source, the 1KGP-LRSC callset contributed the largest total number and largest unique contribution to variant filtering – i.e. how many variants would be unfiltered if a given callset was removed, provided that no other evidence is present. Based on these unique contributions, the performance of our in-house cohort was relatively similar to the 1KGP-LRSC callset that was generated using the same pipeline. Though many SVs were filtered by both, each callset added value; in Tier 1, for example, 1.2% and 0.7% of variants were uniquely filtered by 1KGP-LRSC and the user cohort, respectively. In contrast, we found the contributions of gnomAD-SV variants were negligible, with most being “masked” by variants from other sources. This presumably reflects the limitations of SV detection with short-read sequencing, and we note gnomAD-SV data may be more useful when filtering a short-read SV input callset. These results may inform future users of SVlog about the size and diversity of cohorts that might be required for efficient variant filtering.

Within the ONT dataset, over 4, 500 SVs triggered one or more ‘keeping’ predicates, and these intersected with the unfiltered calls above to produce 10, 88 and 125 SVs prioritised in Tiers 1, 2 and 3, respectively (Figure 4). Because the tiers are not strictly nested, the union across all three tiers comprises 152 unique candidates, which we consider together below. Many of these are low-confidence: 66% (100/152) are genotyped homozygous-reference and 16% (25/152) carry the FILTER=GT tag, which is why the strictest tier is dominated by the artefacts discussed below. Insertions are prevalent in this set: 121 of the 152 variants (80%) are insertions, and the remainder split between 13 deletions, 12 breakends, 3 duplications and 3 inversions. Their sizes span from 4 bp to 2.5 Mb, with a median of 5.8 kb, and roughly half are non-repetitive, the remainder being tandem-repeat or mobile-element events. As expected for a healthy sample, no candidates matched a ClinVar-pathogenic variant at identical, equivalent or related resolution.

Notably, different SVs were prioritised for different reasons. Figure 4 highlights 37 unique predicate signatures – the collection of rules by which a given SV was prioritised – with the number of predicates in each signature ranging from 4 to 11. We found some signatures to appear misleading at first, due to our gene annotations; specifically, a signature may aggregate variants that are both “intronic” and “nonintronic”, which simply implies a combined effect on some overlapping genes that the variant disrupts (e.g. *MMP26* and *OR51A2*/*OR51A4*, *FIRRM* and *SCYL3*, and *ABCC11* and *LONP2*).

The most commonly triggered predicates among prioritised SVs were loss-of-function effects: 138 of the 152 candidates triggered a LoF predicate, most often moderate-confidence LoF (118) and frequently from SVs disrupting intronic regions (87). By contrast, STR predicates — novel STR/VNTR expansions or expansions at known pathogenic STR loci — were comparatively rare, triggered by only 17 candidates. Only 22 candidates overlap a coding sequence, and these cluster in gene families that are known to be structurally polymorphic in healthy populations. Examples include mucins (*MUC3A*, *MUC4*, *MUC6*), the NBPF neuroblastoma-breakpoint cluster (*NBPF12/19/20*), hornerin (*HRNR*), olfactory receptors (*OR2T11*), keratin-associated proteins (*KRTAP10-6*), MAGE genes (*MAGEC1*), and the natural-killer-cell-receptor cluster (*KLRC1*/*KLRC3*). These candidates tend to surface only in Tier 3, driven solely by the permissive `prioritise_lof_any()` predicate, because these structurally polymorphic genes are not sufficiently constrained and so never satisfy the constraint-based predicates that gate Tiers 1-2; moreover, several candidates were low-confidence calls (FILTER=GT or genotyped `0/0`), i.e. false positives. The three largest events illustrate the two reasons extreme calls survive prioritisation in a healthy genome. The largest well-supported call, a 224 kb inversion across an olfactory-receptor cluster on 1q44 (*OR2T11*; genotyped 0|1), lies in a structurally polymorphic region and is only prioritised in Tier 3. The two larger calls — a 2.5 Mb chrX inversion and a 663 kb deletion spanning the 17q21.31 *KANSL1*/*MAPT* locus — are genotyped as homozygous reference (VAF ≤ 0.16, ≤ 8 supporting reads) and are low-confidence calls in segmentally duplicated regions rather than genuine events.

A total of 10 SVs triggered our highest-tier `prioritised1()` predicate, thus warranting consideration. They included a single SV triggering the `prioritise_str_path()` predicate. This was a heterozygous 117 bp insertion in intron 1 of *STARD7*, at the STRchive-catalogued FAME2 locus. However, the inserted repeat does not carry the pathogenic FAME2 motif (the observed motif is TAAAC, not the disease-associated ATTTC), and even its total repeat copy number (33.8) falls far short of the pathogenic range (≥ 274 pathogenic-motif repeats). We therefore interpret this call as an expanded but non-pathogenic allele at this polymorphic, disease-associated STR site rather than a real pathogenic expansion. The remaining nine Tier 1 events comprise the spurious 17q21.31 deletion described above and eight inter-chromosomal breakend (BND) calls in highly constrained genes (*KMT2C*, *LAMA5*, *DOCK9*, *DLGAP4*, *RTN4R*, *MAP3K9*, and *FOXN3* with two adjacent calls). Each of these BNDs is supported by only 5-12 reads at a variant-allele fraction of 0.12-0.57, and five of the eight are genotyped as homozygous reference (0/0), with their mates mapping to disparate telomeric or centromeric regions (features characteristic of low-support Sniffles2 artefacts in complex regions rather than genuine gene-disrupting rearrangements).

We found the results on PacBio data to be generally similar (Figure S3). However, even after accounting for the slight difference in the variant counts, we noticed that there were nearly three times fewer prioritised variants, despite the overall per-predicate rates being comparable (with the only exception of LoF translocations). As expected, for PacBio data, our own large cohort (generated predominantly with PacBio) would come up more often as the reason for filtering variants out, whereas for ONT data, the smaller set of 1KGP-LRSC samples would be more prevalent. Incidentally, fewer signatures could be observed with PacBio data compared to ONT, with the number of completely unfiltered signatures also being around 2 times lower; however, this reduction in the number of signatures only affects the rare, highly specialised signatures.

Overall, SVlog successfully reduced the large SV callsets from LRS data on HG002 to a small and hierarchical list of SVs that could be manually curated. As expected, no notable pathogenic candidate variants were detected in this healthy individual.

### Analysis of patients with rare inherited disease

To further evaluate SVlog, we analysed SV callsets from three patients with rare inherited disease undergoing family-trio PacBio LRS (LRS00037, LRS00038 and LRS00025). Each harboured a disease-causing *de novo* SV, identified previously, providing an opportunity to test SVlog’s prioritisation strategy (Table 1).

**Table 1.**
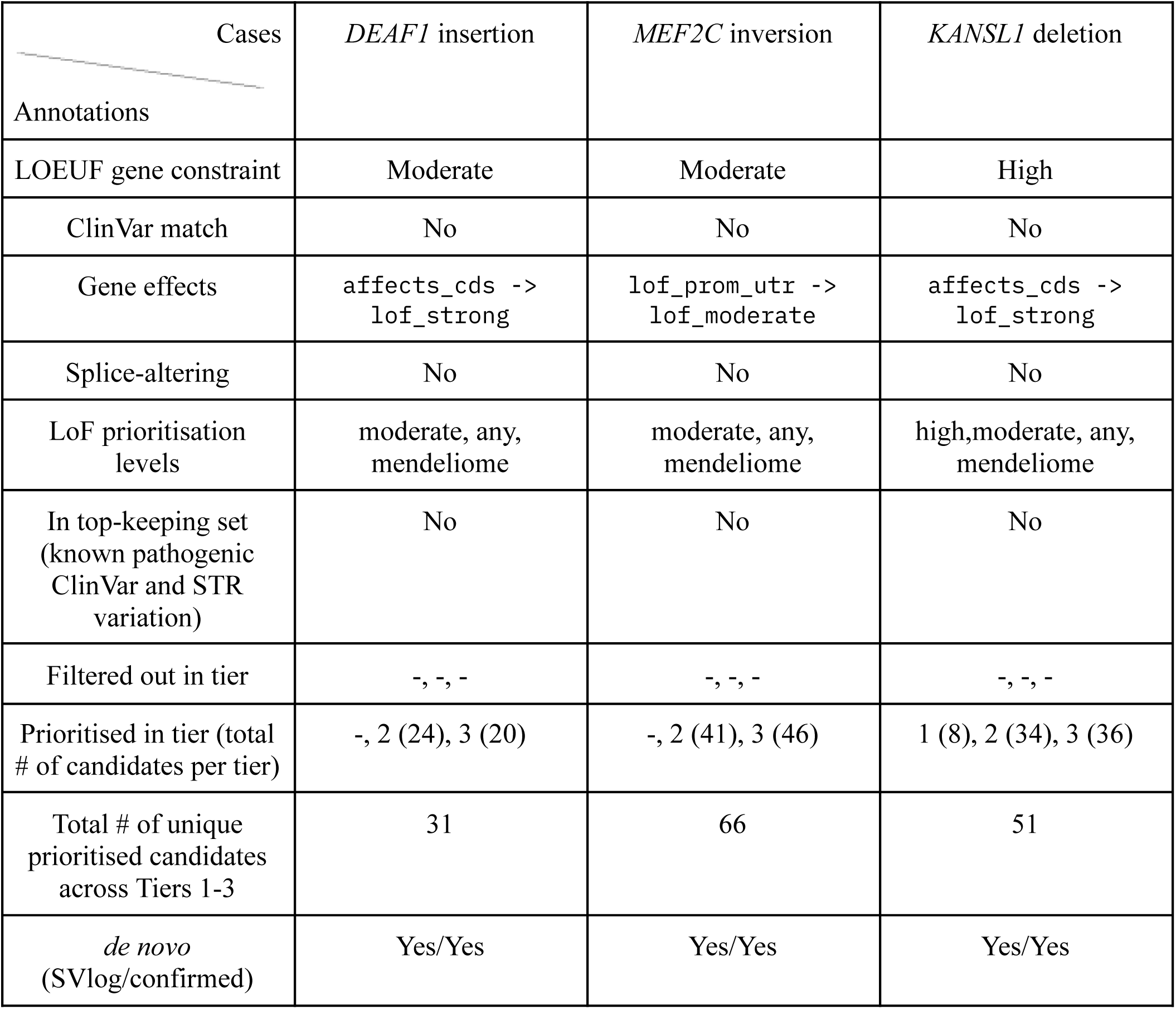
SVlog output for some previously validated clinical cases (PacBio data).

From among many thousands of SVs detected in each proband, SVlog returned 31, 66 and 51 candidates triggering one or more of our prioritisation predicates. Similarly to the HG002 analysis, a substantial fraction (around a third) of these variants were spurious calls. In each case, the known pathogenic SV was among this list of prioritised candidates. Specifically, these were an insertion in the gene *DEAF1*, an inversion impacting *MEF2C* and a deletion in *KANSL1*. These SVs were rare and absent from ClinVar, and all successfully bypassed each of the three tiers of `filtering_out` predicates. Only the *KANSL1* deletion triggered the `prioritise_lof_high()` predicate by disrupting the CDS region of a highly constrained gene, and was prioritised in all three tiers, as a result. The *DEAF1* insertion was also predicted to have a `lof_strong()` gene impact, but the suggested constraint level of the *DEAF1* gene led it to appear only in Tier 2 and Tier 3 prioritised lists. In contrast, the inversion, which does not impact the CDS region of *MEF2C* and instead triggered the `lof_prom_utr()` predicate, was considered only moderately disruptive, and hence also did not make it to the Tier 1 list.

Given the target SV was not the only event prioritised in these three cases, we next considered how a variant curator could further narrow the SVlog-prioritised list down to the causal SV. In each family, the inheritance pattern was consistent with a potential *de novo* variant and a phenotype-relevant gene panel was available (autism, genetic epilepsy and intellectual disability, respectively). Applying SVlog’s `denovo()` predicate to the prioritised SV lists reduced these lists to 9, 19 and 16 candidates. The `recessive()` predicate was also evaluated for each proband, but it yielded no suitable candidates, so the *de novo* route was used throughout. Subsequently restricting these to the appropriate gene panel via the `prioritise_lof_my_panel()` predicate isolated the known causal SV as a single unambiguous hit in each case. The remaining *de novo* candidates fall into three broad classes and are readily de-prioritised: intergenic or fully intronic events unlikely to disrupt a coding gene; variants affecting genes not on the phenotype-relevant panel (for example, *SIGLEC14* in LRS00037, *TOGARAM2* and *SLC9A3* in LRS00038, and *LHFPL3* in LRS00025); and low-support inter-chromosomal breakend (BND) calls that recur at identical coordinates across independent samples, most strikingly chr7:152416667 (annotated as *KMT2C*), which appears as a *de novo* BND in all three clinical cases and is also called in HG002, and is therefore readily identified as a systematic Sniffles2 artefact on IGV inspection.

In summary, SVlog prioritised known pathogenic SVs among a small number of candidates produced via agnostic analysis. When combined with basic phenotypic information and consideration of the inheritance pattern, the known pathogenic SV remained as a single prioritised event in each patient.

## Discussion

The desire to delegate medical decision-making to a “black box” comes at a cost: AI models, although useful for basic science, typically struggle to meet the transparency and reliability standards expected of medical tools (Quinn *et al*., 2022). We believe that, where possible, a simple deterministic model should be prioritised over a more complete yet stochastic one.

Structural variants remain challenging to annotate, filter and prioritise. Even though tools like AnnotSV and SVAnnotate have been available for years, they do not provide a direct way to edit and expand their annotation capabilities (Geoffroy *et al*., 2018; Collins *et al*., 2020). In contrast, with our meta-programming framework, we were able to re-implement some annotations from those tools, such as the “copy gain” prediction, and enhance their functionality.

With SVlog, we defined a prototype of a deterministic model for variant prioritisation in the context of disease studies, in the form of a computer library. This model can be scrutinised directly from the source code of the library. We defined a set of possible relations between structural variants as rules using the logic programming paradigm. We opted for Soufflé Datalog as our modelling tool (Jordan *et al*., 2016). When the implemented rules are used through the SVlog library, the Soufflé engine can derive new facts about input data by applying those rules.

Our library is perhaps most similar to the ‘bio_db’ library developed in Prolog for the integration of various biological databases (Angelopoulos and Giamas, 2015). Even though SVlog could be categorised as a pipeline by some, we would like to emphasise that in traditional pipelines (e.g. Snakemake, WDL) the end goal is the derivation of some final piece of data through intermediate stages, in a step-by-step manner. Here, however, there is no end goal *per se*. Workflows provide a way to decompose a task into parts, but the data itself is still processed at a low level of abstraction. Instead, here the user can just pull predicates from the library and request to see output for those predicates after providing input data, all at a much higher level of abstraction. Therefore, there is no process or workflow to automate within this framework, and that is why it is fundamentally different to other common computational approaches in bioinformatics.

Importantly, our SVlog system can be easily adapted to accommodate various additional databases and genome reference data. For example, the hg38-to-chm13 liftover procedure for genomic elements is as simple as updating their loci data in the relevant predicates. Furthermore, the functionality employed in SVlog is not limited to Datalog’s standard operators, as Soufflé can load functions implemented externally in C/C++ (also known as ‘functors’) and use them in the processing of the rules, which can be particularly useful in bioinformatics for string manipulation and other use cases.

### Limitations and Future Work

Despite the simplicity and minimalism of Datalog, some aspects of Soufflé must be considered when working with and expanding our framework. In particular, the Magic Set Transformations that our library relies on cause some structurally sound rules to fail, because of current limitations of Soufflé. Specifically, as of yet, aggregations are not supported when applying Magic Sets in Soufflé. As a result, we had to implement some rules as negations of positive examples. Although such implementations may not look intuitive or elegant, our library serves as a prototype that will hopefully be developed further with continuous improvements made to Datalog systems.

We are also interested in incorporating other logic programming tools in our system, such as Clingo ASP (Gebser *et al*., 2019) and Scallop (Li *et al*., 2023). This would enable us to extend the scope of SVlog to more complex search problems within the variant prioritisation domain.

### Conclusion

The logic programming framework presented here provides a fast and efficient way of working with structural variants, both in terms of development and performance. The Datalog language used here makes the developed library easily maintainable, highly extensible, transparent and reliable, enabling interpretable and reproducible analyses. Our approach addresses the computational challenges associated with the processing of structural variants, at a high level of abstraction and in a manner that is algorithm-free. We hope that this novel framework will spark new ideas in terms of collaborative efforts to summarise current knowledge and best practices with regard to genetic variation in the medical community.

## Code availability

SVlog is publicly available at https://github.com/GenTechGp/SVlog

