## Supplementary material for "SVlog: a logic programming framework for understanding structural variation in genomic disease"

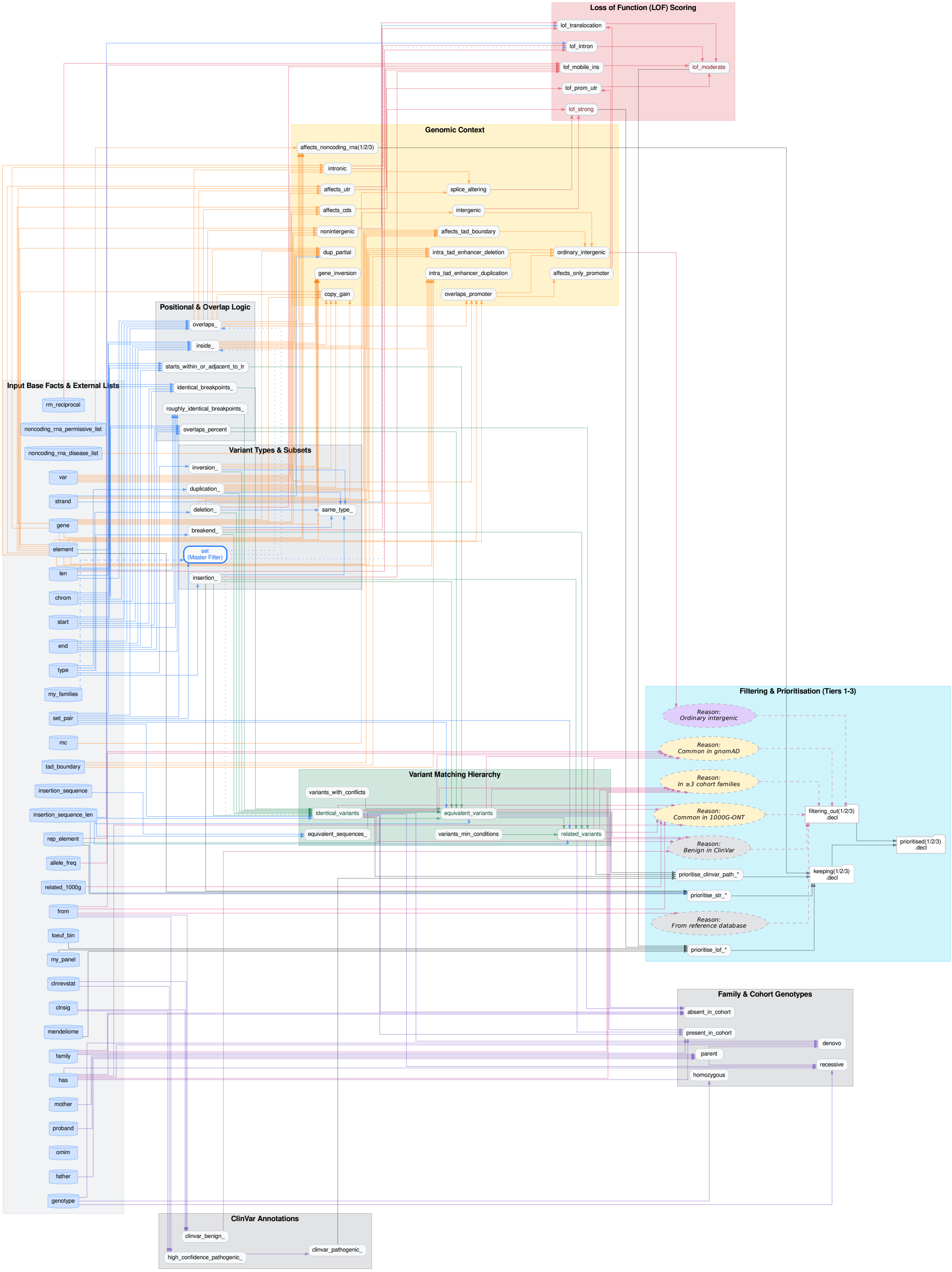


Figure S1. Complete flow of input and output predicates in SVlog (function calls).

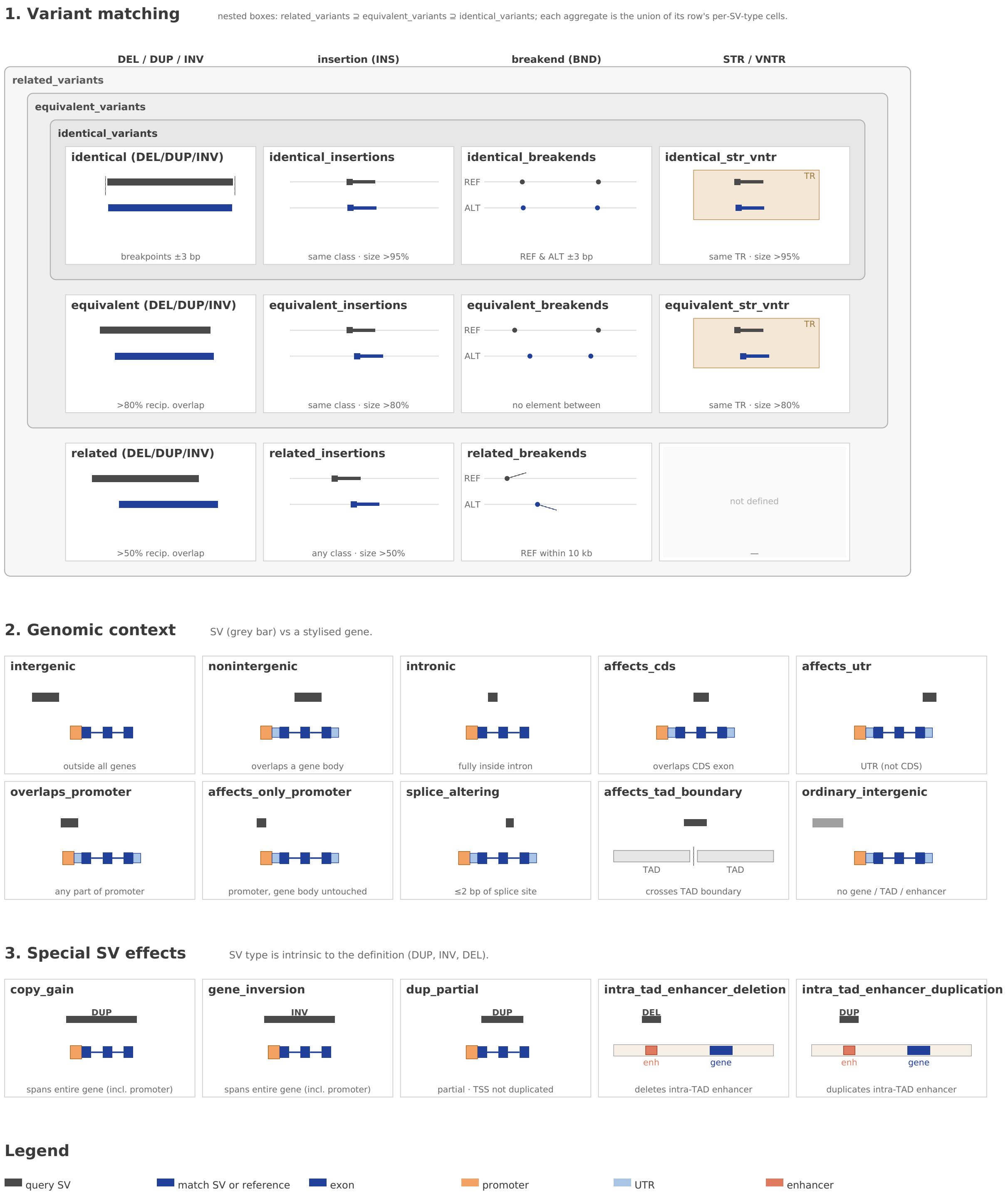


Figure S2. Visual definitions of key variant-matching and annotation predicates in SVlog.

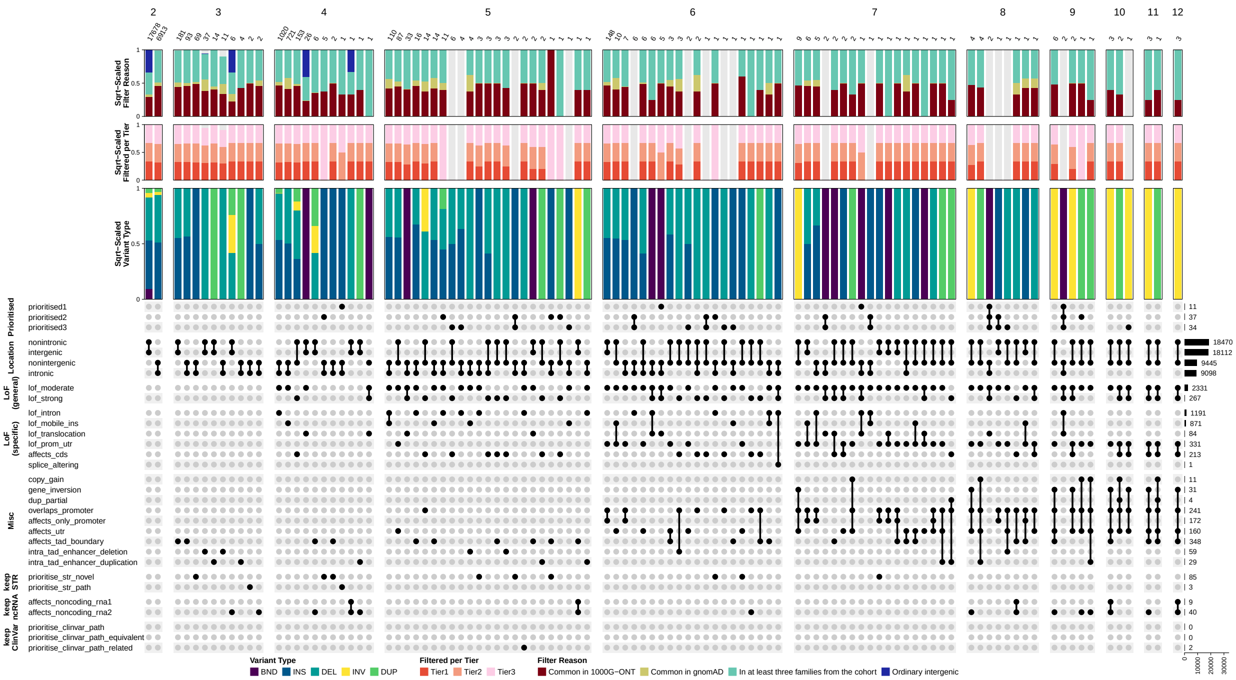


Figure S3. Prioritisation and variant effect predicates (rows) and predicate signatures (columns). HG002-PacBio data (27,557 variants). Only the most relevant predicates are included. Selectivity of each predicate is shown on the right. The panes are split based on the number of predicates in each signature.

Table S1. Key SVlog predicates.

| Predicate name | Group | Description | Comments | VCF INFO |
|---|---|---|---|---|
| identical_insertions | 1 | Two SVs have INS type; AND<br>Same SV classification; AND<br>Left breakpoints within ∓ 3bp; AND<br>Length of smaller INS is >95% of larger INS; | | · |
| identical_str_vntr | 1 | Two SVs are INS-INS or DEL-DEL; AND<br>SV classification is ‘tandem repeat’ with STR or VNTR subclass (for both); AND<br>Left breakpoints are within or adjacent to (within 50bp) the same genomic tandem repeat element; AND<br>Length of smaller SV is >95% of larger SV | *Filtering against* identical_str_vntr *should identify expansions or contractions with unique size or repeat motif, compared to common alleles at a given repeat site* | · |
| identical_breakends | 1 | Two SVs have BND type; AND<br>the two REFs are within ∓ 3 bp; AND<br>the two ALTs are within ∓ 3 bp | | · |
| identical_variants | 1 | identical_insertions; OR<br>identical_str_vntr; OR<br>identical_breakends; OR<br>Two SVs have same SV type; AND<br>SV type is DEL, DUP or INV; AND<br>Share two breakpoints that are both within ∓ 3 bp | Used for filter_identical (filtering_out1) | · |
| equivalent_insertions | 1 | Two SVs have INS type; AND<br>Same SV classification; AND<br>Distance between their left breakpoints is <20% length of smaller INS; AND<br>Length of smaller INS is >80% of larger INS | should include identical_insertions by design | · |
| equivalent_str_vntr | 1 | Two SVs are INS-INS or DEL-DEL; AND<br>SV classification is ‘tandem repeat’ with STR or VNTR subclass (for both); AND<br>Left breakpoints are within or adjacent to (within 50bp) the same genomic tandem repeat element; AND<br>Length of smaller SV is >80% of larger SV | *Filtering against* equivalent_str_vntr *should identify significant outliers in size compared to common alleles at a given repeat site* | · |

| | | | | |
|---|---|---|---|---|
| equivalent_breakends | 1 | Two SVs have BND type; AND<br>their REFs as well as their ALTs are within their respective *element-level* neighbourhoods (i.e. there are no important genomic elements that start and/or end between the two BNDs - this is similar to the checks in equivalent_variants). | should include identical_breakends by design | · |
| equivalent_variants | 1 | identical_variants; OR<br>equivalent_insertions; OR<br>equivalent_str_vntr; OR<br>equivalent_breakends; OR<br>"Two SVs have same SV type; AND<br>SV type is DEL, DUP or INV; AND<br>Share genome coordinates with >80% reciprocal overlap; AND<br>Have no conflicts in gene impact prediction (structural effects)." | *Conflicting structural effects arise whenever there is an "event" taking place inside either one of such sequences. If there is an important genomic element which has either its START or END inside either of the two sequences, that constitutes an "event". If the element starts inside the sequence, we don't care where it ends; if it ends inside the sequence, we don't care where it starts. That is, there is no event if and only if the sequence is entirely inside (contained by) the element. In the most general sense, "elements" constitute everything that is not a variant.*<br><br>Used for filter_equivalent (filtering_out2) | · |
| related_insertions | 1 | Two SVs have INS type; AND<br>Distance between their left breakpoints <50% length of smaller INS; AND<br>Length of smaller INS is >50% of larger INS | should include equivalent_insertions and identical_insertions by design | · |
| related_breakends | 1 | Two SVs have BND type; AND<br>Their REF positions are within a 10000bp window (regardless of where their ALTs are pointing); | (should include equivalent_breakends and identical_breakends) | · |
| related_variants | 1 | equivalent_variants; OR<br>related_insertions; OR<br>related_breakends; OR<br>"Two SVs have same SV type; AND<br>SV type is DEL, DUP or INV; AND<br>Share genome coordinates with >50% reciprocal overlap" | Used for filter_related (filtering_out3) | · |
| common_gnomad * | 2 | SV in gnomAD with AF > 0.01 | | . |

| | | | | |
|---|---|---|---|---|
| common_1kg * | 2 | SV in 1KG-ONT in >=3 individuals | | . |
| common_internal * | 2 | SV in our internal study cohort in >=3 families | | . |
| clinvar_benign # | 2 | SV from ClinVar with classification of ‘Benign’ - criteria provided, multiple submitters, no conflicts (*high confidence*) | | . |
| clinvar_pathogenic # | 2 | SV from ClinVar with classification of ‘P’, ‘LP’, ‘P/LP’ - reviewed by expert panel, single submitter (with criteria provided), or multiple submitters (with criteria provided and no conflicts in interpretation) (*medium confidence*) | | . |
| clinvar_vus # | 2 | SV from ClinVar with classification of ‘Uncertain significance’ | | . |
| intergenic | 3 | SV is *not* nonintergenic, i.e. does not overlap any genes ("transcript" elements, which do not include the promoter region) | | intg |
| nonintergenic | 3 | NOT intergenic | | nonintg |
| ordinary_intergenic | 3 | SV does not overlap any protein coding genes (hence NOT affects_cds) or promoters; AND<br>is NOT affects_tad_boundary; AND<br>does not delete or duplicate any intra-TAD enhancers | | . |
| affects_cds | 3 | SV affects (i.e. overlaps) any part of a CDS region. Just like with all other affects/overlaps predicates, the overlaps can be produced by any SV (deletions, insertions, inversions, duplications, BNDs). Variants with "copy_gain" or "gene_inversion" effects are *not* included. | | affects_cds |
| splice_altering | 3 | SV is intronic (hence NOT affects_cds) and affects a splice donor or splice acceptor site (2 bp before/after an exon) | | splice_altering |
| intronic | 3 | SV is fully within an intron (hence NOT affects_cds) | | intr |
| affects_utr | 3 | SV affects (i.e. overlaps) a UTR region; AND<br>NOT affects_cds | such a variant will affect one of the UTRs - and also possibly promoter/enhancers - but nothing within the ORF | affects_utr |

| | | | | |
|---|---|---|---|---|
| overlaps_promoter | 3 | SV overlaps the promoter of a gene | | . |
| affects_only_promoter | 3 | SV affects (i.e. overlaps) the promoter of a gene, but not the gene itself ("transcript" element) | (note: enhancers or other elements related to the gene may also be affected) | affects_only_promoter |
| affects_tad_boundary | 3 | SV affects (i.e. overlaps) a TAD boundary (region between two TADs that are within 100kb from each other; if they are more than 100kb apart, the boundary is considered "highly extended", which there are only ~30% of). | https://doi.org/10.1073/pnas.2413112122 | affects_tad_boundary |
| affects_noncoding_rna1 | 3 | Affects (i.e. overlaps) a very important ncRNA gene (seqr known disease snRNA list) | | affects_noncoding_rna1 |
| affects_noncoding_rna2 | 3 | Affects (i.e. overlaps) a somewhat important ncRNA gene (seqr permissive snRNA list) | | affects_noncoding_rna2 |
| affects_noncoding_rna3 | 3 | Affects (i.e. overlaps) *some* ncRNA gene (any ncRNA gene from GENCODE) | | affects_noncoding_rna3 |
| lof_strong | 3 | Query SV is nonintergenic; AND<br>affects_cds or splice_altering or lof_translocation (i.e. has to be within a gene) | | lof_strong |
| lof_moderate | 3 | Query SV is lof_prom_utr or lof_intron or lof_mobile_ins or lof_translocation (i.e. can be within a gene or next to it) | | lof_moderate |
| lof_prom_utr | 3 | Query SV affects_only_promoter or affects_utr in a gene | | lof_prom_utr |
| lof_intron | 3 | Query SV is intronic; AND<br>"SV length is >50% of the length of the intron size it is contained within; OR<br>The distance to the nearest splice site is <500bp" | | lof_intron |
| lof_mobile_ins | 3 | Query SV has INS or DEL type; AND<br>Query SV is intronic OR affects_utr OR affects_only_promoter; AND<br>SV classification is 'mobile element' (i.e. "DNA", "LINE", "SINE", "LTR" and "Retroposon" types) with ‘transposition’ status (RM_RECIPROCAL=Full) | | lof_mobile_ins |
| lof_translocation | 3 | Query SV has BND type; AND<br>REF position is anywhere within a gene (i.e. nonintergenic or affects_only_promoter); OR<br>REF position is within 25kb of the TSS | | lof_translocation |

| | | | | |
|---|---|---|---|---|
| | | (strand-dependent) | | |
| copy_gain | 3 | Query SV has a copy-gain effect on a gene. | This is a re-implementation of the "PREDICTED_COPY_GAIN" annotation from SVAnnotate: *"Gene(s) on which the SV is predicted to have a copy-gain effect. This occurs when a duplication spans the entire transcript, from the first base of the 5' UTR to the last base of the 3' UTR."* | copy_gain |
| gene_inversion | 3 | Query SV has a gene-inversion effect on a gene. | This is a re-implementation of the "PREDICTED_INV_SPAN" annotation from SVAnnotate: *"Gene(s) which are entirely spanned by an SV's inversion. A whole-gene inversion occurs when an inversion spans the entire transcript, from the first base of the 5' UTR to the last base of the 3' UTR."* | gene_inversion |
| dup_partial | 3 | This is a re-implementation of the "PREDICTED_DUP_PARTIAL" annotation from SVAnnotate: *"Gene(s) which are partially overlapped by an SV's duplication, but the transcription start site is not duplicated. The partial duplication occurs when a duplication has one breakpoint within the transcript and one breakpoint after the end of the transcript. When the duplication is in tandem, the result is that there is one intact copy of the full endogenous gene."* | | dup_partial |
| genes_affected | 3 | If query SV affects at least one element that's associated with a gene, that gene is considered to by affected by the SV (for enhancers, if the associated gene information is available, it's sufficient for an SV to affect just that enhancer for it to "affect" the corresponding gene) | | . |
| in | | SV is in some database (either *comes* from that database or has an identical_variants SV in that database) | | . |
| only_in | | SV is only in that one database (e.g. only in 1000G or only in gnomAD) | | . |
| filtering_out1 | 4 | Query SV is ordinary_intergenic; OR<br>When compared using identical_variants the query matches an SV that is common_gnomad; OR | filter_identical *provides strong evidence that the query SV is not pathogenic, because an identical SV(s) that is* | fo1 |

| | | | | |
|---|---|---|---|---|
| | | common_1kg; OR<br>common_internal; OR<br>clinvar_benign | *common/benign was identified (tier 1)* | |
| filtering_out2 | 4 | Query SV is ordinary_intergenic; OR<br>When compared using equivalent_variants the query matches an SV that is common_gnomad; OR<br>common_1kg; OR<br>common_internal; OR<br>clinvar_benign | filter_equivalent *provides strong+moderate evidence that the query SV is not pathogenic, because an equivalent SV(s) that is common/benign was identified (tier 2)* | fo2 |
| filtering_out3 | 4 | Query SV is ordinary_intergenic; OR<br>When compared using related_variants the query matches an SV that is common_gnomad; OR<br>common_1kg; OR<br>common_internal; OR<br>clinvar_benign | filter_related *provides strong+moderate+some evidence that the query SV is not pathogenic, because a related SV(s) that is common/benign was identified (tier 3)* | fo3 |
| prioritise_clinvar_path | 5 | When compared using identical_variants the query matches a variant that is clinvar_pathogenic | | prioritise_clinvar_path |
| prioritise_clinvar_path_equivalent | 5 | When compared using equivalent_variants the query matches a variant that is clinvar_pathogenic | | prioritise_clinvar_path_equivalent |
| prioritise_clinvar_path_related | 5 | When compared using related_variants the query matches a variant that is clinvar_pathogenic | | prioritise_clinvar_path_related |
| prioritise_lof_high | 5 | Query SV is lof_strong in a gene that is highly constrained (LOEUF top 20%) | | prioritise_lof_high |
| prioritise_lof_mod | 5 | Query SV is lof_moderate or lof_strong in a gene that is moderately constrained (LOEUF top 50%) | | prioritise_lof_mod |
| prioritise_lof_mendeliome | 5 | Query SV is lof_moderate or lof_strong in a gene that is green on the mendeliome panel in panelApp | | prioritise_lof_mendeliome |
| prioritise_lof_my_panel | 5 | Query SV is lof_moderate or lof_strong in a gene on the <user provided panel> in panelApp | | prioritise_lof_my_panel |
| prioritise_lof_any | 5 | Query SV is lof_moderate or lof_strong in any gene | | prioritise_lof_any |
| prioritise_str_path | 5 | Query SV type is INS; AND | | prioritise_str_path |

| | | | | |
|---|---|---|---|---|
| | | SV classification is ‘tandem repeat’ with STR or VNTR subclass; AND<br>Left breakpoint is within or adjacent to (within 50bp) a genomic tandem repeat element that is a known pathogenic STR site; AND<br>SV length is >50% size of the underlying STR site | | |
| prioritise_str_novel | 5 | Query SV type is INS; AND<br>SV classification is ‘tandem repeat’ with STR or VNTR subclass; AND<br>Query SV is within a gene (any part of the gene including the promoter and UTR - i.e. nonintergenic OR affects_only_promoter); AND<br>does **not** have 3 equivalent_str_vntr variants, each with a length greater than the query SV; AND<br>SV length is >50% larger than the *genomic tandem repeat* element the SV breakpoint is within or adjacent to (within 50bp) | | prioritise_str_novel |
| absent_in_cohort1 | 5 | Query SV has no identical variants in other families in the cohort | | . |
| absent_in_cohort2 | 5 | Query SV has no equivalent variants in other families in the cohort | | . |
| absent_in_cohort3 | 5 | Query SV has no related variants in other families in the cohort | | . |
| absent_in_1000g1 | 5 | Query SV has no identical variants in 1000G | | . |
| absent_in_1000g2 | 5 | Query SV has no equivalent variants in 1000G | | . |
| absent_in_1000g3 | 5 | Query SV has no related variants in 1000G | | . |
| homozygous | 7 | SV has genotype "1/1" | | . |
| denovo | 7 | Proband's SV has no identical_variants among their parents' SVs, *but data from both parents is available* | | denovo |
| possible_denovo | 7 | like denovo but not denovo because only one parent is available | | possible_denovo |
| recessive | 7 | Proband's SV is<br>homozygous, AND<br>identical_variants with an SV from their mother, AND<br>identical_variants with an SV from their father. | | recessive |
| keeping0 | 8 | prioritise_clinvar_path OR prioritise_str_path | | keeping0 |

| | | | | |
|---|---|---|---|---|
| keeping1 | 8 | prioritise_clinvar_path<br>OR<br>prioritise_lof_high<br>OR<br>prioritise_str_path<br>OR<br>prioritise_lof_my_panel<br>OR<br>affects_noncoding_rna1 | prioritise_tier1 (Strong evidence of pathogenicity) | keeping1 |
| keeping2 | 8 | prioritise_clinvar_path_equivalent<br>OR<br>prioritise_lof_mod<br>OR<br>prioritise_lof_mendeliome<br>OR<br>prioritise_str_path<br>OR<br>prioritise_str_novel<br>OR<br>affects_noncoding_rna2 | prioritise_tier2 (Moderate evidence of pathogenicity) | keeping2 |
| keeping3 | 8 | prioritise_clinvar_path_related<br>OR<br>prioritise_lof_any<br>OR<br>prioritise_str_path<br>OR<br>prioritise_str_novel<br>OR<br>affects_noncoding_rna2 | prioritise_tier3 (Some evidence of pathogenicity) | keeping3 |
| prioritised1 | 8 | keeping1 AND NOT filtering_out1 | tier 1 prioritisation | prioritised1 |
| prioritised2 | 8 | keeping2 AND NOT filtering_out2 | tier 2 prioritisation | prioritised2 |
| prioritised3 | 8 | keeping3 AND NOT filtering_out3 | tier 3 prioritisation | prioritised3 |

Predicates marked with * are virtual/nonexistent, which means they are only implemented as Soufflé code and can be re-used only by copying the code (because of performance considerations).
Predicates marked with # are auxiliary predicates used for other predicates - they cannot be evaluated on the target set of SVs directly.
